# Twin studies with unmet assumptions are biased towards genetic heritability

**DOI:** 10.1101/2020.08.27.270801

**Authors:** Edwin S. Dalmaijer

## Abstract

For a century [1,2], studies of monozygotic and dizygotic twins have yielded estimates of trait heritability. The clever logic behind them is that while both types of twins share environments, their genetic overlap is different. Hence, larger trait correlations between monozygotic compared to dizygotic twins indicate heritability (nature), whereas similar correlations indicate shared environmental influences (nurture), and low correlations indicate shaping through non-shared environments (external influences and measurement error). While many have written on the assumptions that both types of twins share equal environments [3–5], and that parental genetics and environment are independent [6,7]; fewer have put their data where their mouth is. Here, the impacts of unmet assumptions were investigated using a generative mixture model of twin phenotypes. The results indicated that violations of the equal environments assumption yielded large overestimations of heritability and underestimations of shared environmental influences. On the other hand, when parental genetics shaped twins’ shared environments, only minor non-linear biases against heritability emerged. Finally, realistic levels of measurement error uniformly depressed estimates for genetic and shared environmental factors. In sum, twin studies are particularly susceptible to overestimation of genetic and non-shared environmental influences. This bias could explain why some traits, such as attitudes towards property taxes [8], show suspiciously high heritability without a biologically plausible mechanism. Particularly in the context of traits with convincing mechanisms of cultural transmission [9–11] and complex gene-environment interactions [6], researchers should not allow biases in twin studies to overestimate heritability.

## Results

Three of the main potential causes of bias in twin studies were simulated. First, the equal environments assumption was modelled as direct statistical dependencies between twins’ genetics and their shared environments, non-shared environments, or both. Second, parental genetic confounding was modelled as statistical dependency between parental genotype and twins’ shared environment. Finally, measurement error was introduced as normally distributed noise on the trait.

### Simulated parameter recovery

Simulated parameters were accurately recovered using traditional methods (Figure S2) and structural equation modelling (SEM). In the absence of any simulated confounds, SEM underestimated heritability by 1.2 percentage points [*M*=-0.012, *SD*=0.093, *t*(5785)=-9.42, *p*<0.001, *d*=-0.12], and overestimated shared [*M*=0.006, *SD*=0.086, *t*(5785)=5.29, *p*<0.001, *d*=0.07] and nonshared environment [*M*=0.006, *SD*=0.026, *t*(5785)=16.32, *p*<0.001, *d*=0.21] by 0.6 percentage points each. These results indicate that there were very small but statistically detectable biases in either the generative model or in twin methodology.

### Equal environment assumption

Statistical methods in twin studies assume that monozygotic and dizygotic twins are reared in environments that are equally similar for both. This assumption is violated if children’s behaviour or appearance inspires differential treatment (“evocative genetic influence”, [12]), or if they help shape their environment or experience thereof (“active genetic influence”, [12]). In other words, an individual’s genes could confound their environment, and this would create more similar environments for monozygotic than dizygotic twins.

Even at only 10% confounding through gene-environment interaction, heritability was overestimated by 23.1 percentage points [*M*=0.231, *SD*=0.136, *t*(5807)=129.69, *p*<0.001, *d*=1.70], while shared environment was underestimated by 9.8 points [*M*=-0.098, *SD*=0.131, *t*(5807)=-56.63, *p*<0.001, *d*=-0.74], and non-shared by 13.3 points [*M*=-0.133, *SD*=0.085, *t*(5807)=-119.18, *p*<0.001, *d*=-1.56]. Even when counting the 10% confound as “genetic” influence, heritability was overestimated by 16.2 percentage points [*M*=0.162, *SD*=0.127, *t*(5807)=69.65, *p*<0.001, *d*=1.27], while shared environment was underestimated by 6.6 points [*M*=-0.066, *SD*=0.115, *t*(5807)=-43.97, *p*<0.001, *d*=-0.58], and non-shared by 9.5 points [*M*=-0.095, *SD*=0.069, *t*(5807)=-105.13, *p*<0.001, *d*=-1.38]. Results for further levels of confounding are reported in Figure 1 (yellow lines), as are results for interactions with only shared (red lines) or non-shared environments (orange lines). As an illustration, misestimations for different simulation conditions at 10% confounding are reported in Figures 2 (both), S3 (shared) and S4 (non-shared environment).

**Figure 1.**
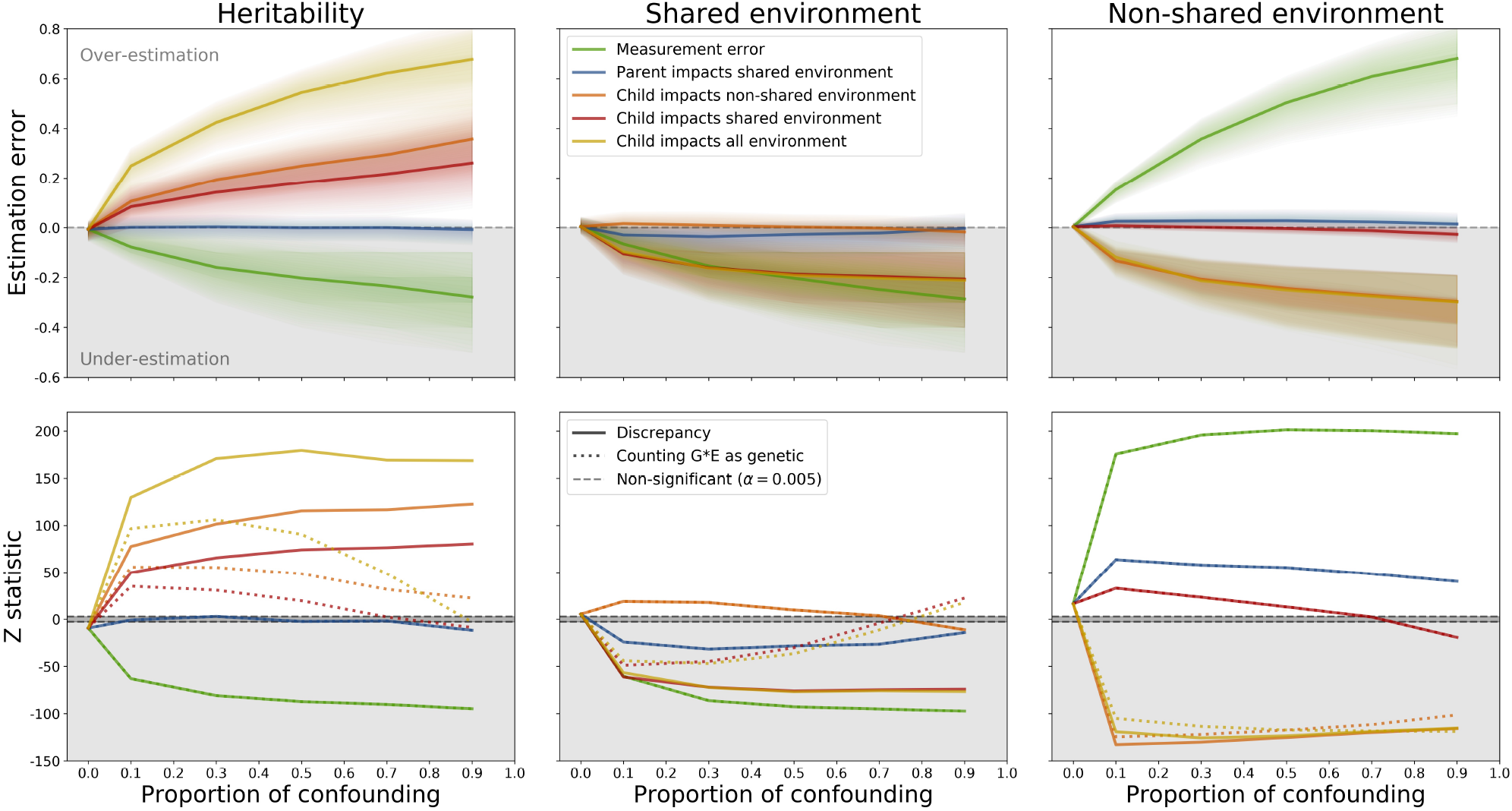
The top rows show median estimation error across a variety of simulations, computed as the difference between ground truth and estimates from structural equation modelling. The shaded areas illustrate the interquartile ranges. Simulations introduced statistical confounds between twin genetics and both environmental components (yellow), or only shared (red), and non-shared environmental influences (orange); or between parental genetics and shared environment (blue); or introduced measurement error (green). In the bottom row, straight lines reflect Z scores for differences between estimates and ground truth, and the dotted line for differences between estimates and ground truth when gene-environment interactions are counted towards ground truth heritability.

**Figure 2.**
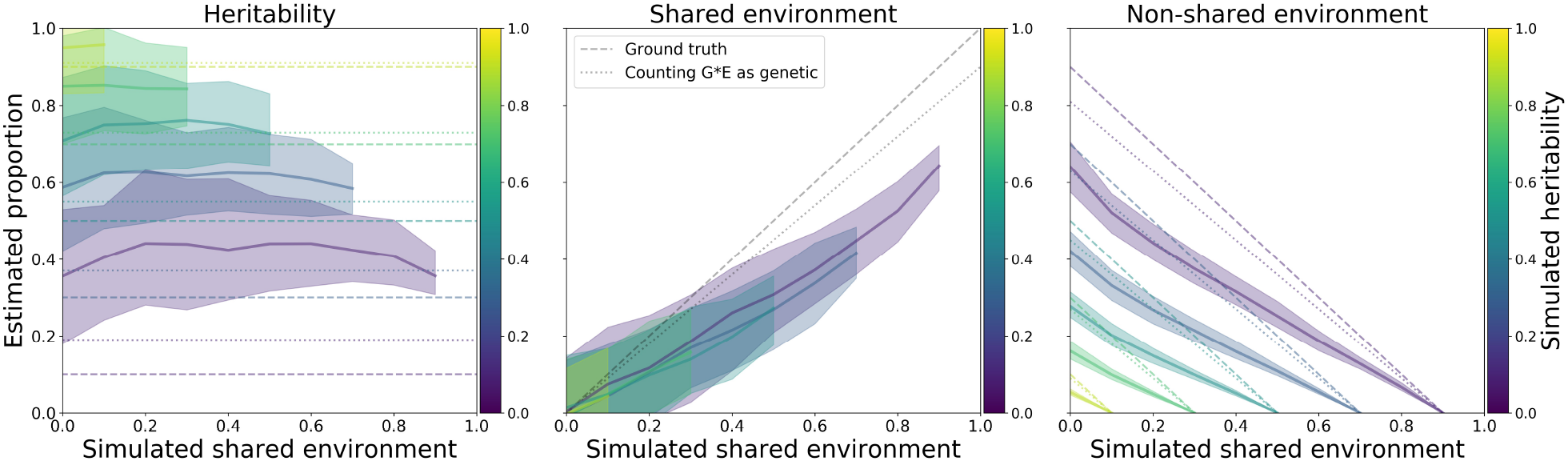
Results from 100 simulations per combination of ground truth proportions in which shared and non-shared environments were 10% confounded by twin genetics. Average estimated heritability (left panel), shared (middle panel), and non-shared environmental influences (right panel). The shaded areas indicate the 5^th^ to 95^th^ percentiles. Dashed lines show the ground truth, and dotted lines ground truth if gene-environment interactions are counted towards heritability.

These results indicate that heritability was not only overestimated when gene-environment interactions were present, but that it was overestimated even when the entire confound was counted as “genetic” influence. Twin methodology thus deviated from the ground truth in the presence of gene-environment interactions.

### Parental genes shape twins’ environment

Twins do not only inherit their parents’ genes, but also grow up in environments that are partly shaped by them. Parents’ genetics can thus confound twins’ shared environment (“passive genetic influence”, [12]). It has been argued that this can result in an overestimation of environmental and an underestimation of genetic influences [7].

This assumption did not uniformly hold true. When 30% of the shared environment was determined by parental genotype (Figure 3), SEM overestimated the heritability by 0.4 percentage points [*M*=0.004, *SD*=0.098, *t*(5793)=2.89, *p*=0.004, *d*=0.04], while shared environment was underestimated by 4.8 points [*M*=-0.048, *SD*=0.114, *t*(5793)=-31.80, *p*<0.001, *d*=-0.42], and nonshared was overestimated by 4.4 points [*M*=0.044, *SD*=0.058, *t*(5793)=57.78, *p*<0.001, *d*=0.76]. Further results are reported in Figure 1 (blue lines).

**Figure 3.**
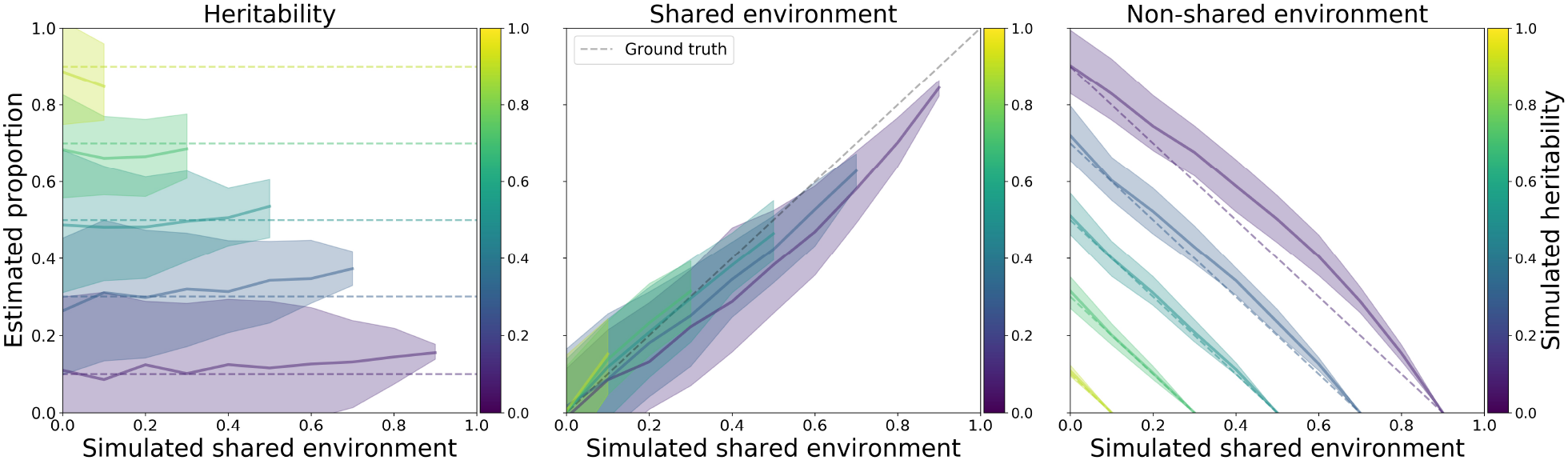
Results from 100 simulations per combination of ground truth proportions in which shared environment was 30% confounded by parental genetics. Average estimated heritability (left panel), shared (middle panel), and non-shared environmental influences (right panel). The shaded areas indicate the 5^th^ to 95^th^ percentiles. Dashed lines show the ground truth.

These results indicate that twin methodology underestimated shared environmental effects and overestimated non-shared environmental effects when parental genetics partially shaped the shared environment. However, these effects on heritability were marginal, and not always in the same direction (overestimation at 30% confounding, but underestimation at 90%).

### Measurement error

It is generally assumed that measurement error is captured by the proportion of variance that is attributed to influences from non-shared environments (e.g. [13]).

This held true: at 30% measurement error (Figure 4), heritability was underestimated by 16.8 percentage points [*M*=-0.168, *SD*=0.169, *t*(6599)=-80.97, *p*<0.001, *d*=-1.00] and shared environment was by 16.2 points [*M*=-0.162, *SD*=0.1524, *t*(6599)=-86.31, *p*<0.001, *d*=-1.06], while non-shared environment was overestimated by 33.0 points [*M*=0.330, *SD*=0.137, *t*(6599)=195.71, *p*<0.001, *d*=2.41]. Further results are reported in Figure 1 (green lines).

**Figure 4.**
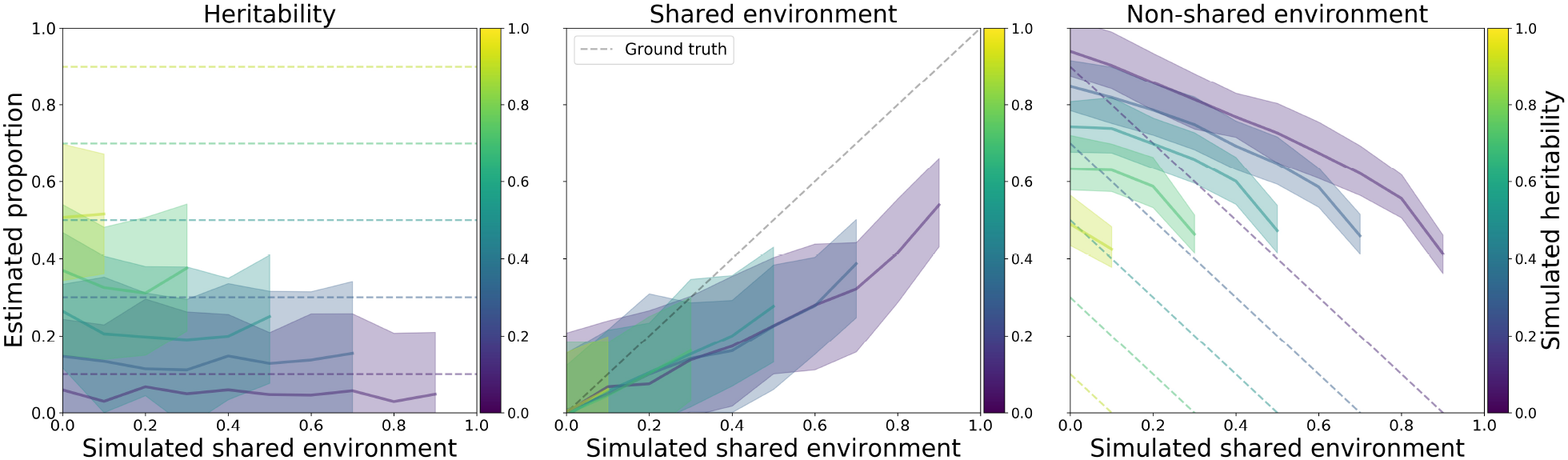
Results from 100 simulations per combination of ground truth proportions in which nonshared environment was 30% confounded by measurement error. Average estimated heritability (left panel), shared (middle panel), and non-shared environmental influences (right panel). The shaded areas indicate the 5^th^ to 95^th^ percentiles. Dashed lines show the ground truth.

### Practical example

To illustrate how the above biases impact studies, the estimates from a recent study on the heritability of disgust proneness [14] are compared to the outcomes of simulations along the environmental confounds spectrum. The study employed an instrument with reliability α=0.79, so measurement error was set to 20%, while the proportion of environmental variance explained by twin and parental genetic confounds was varied between 0 and 1. A total of 2.6% of all simulations produced values within the confidence intervals of the original study (heritability 27-41%, nonshared environment 59-72%). However, ground truths in these simulations included up to 60% heritability, and 30-80% non-shared environmental influences. Crucially, the simulations that produced estimates within the study’s confidence intervals also included ground truths with up to 50% shared environmental influences, whereas the original authors reported none.

These results indicate that when the ground truth gene-environment interaction is unknown, a wide range of true values can produce the same estimates of heritability and environmental influences.

## Discussion

This study sought to quantify the effects of violated assumptions in the decompositional analysis of trait correlations between monozygotic and dizygotic twin pairs, using simulated populations of parents and their offspring. Parental genetic confounds did not impact estimations much, whereas measurement error increased estimated non-shared environmental influences at the cost of genetic and shared environmental influences. Crucially, the current results show that heritability is overestimated when twin’s genetics and environment are confounded, even when this gene-environment interaction was counted towards the ground truth heritability.

This is nowhere near the first work to warn about the pitfalls of twin studies. Past criticism has ranged from thoughtful and precise warnings about specific confounds [3,4,6,7,12], to outright dismissal of twin studies as “pseudoscience” [5]. There have even been attempts to quantify violations of the equal environments assumption in real samples [15], aided by simulations that showed large overestimation of additive genetic effects and underestimation of shared environmental effects. What the current work adds is a wider range of tested assumptions and the effects of their violation, and the surprising finding that heritability is overestimated even if child-environment interactions are fully considered to be “genetic” influences.

Especially early in life, twin’s shared environments are shaped primarily by their parents. Simulations showed that confounding environment and parental genetics only produced minor estimation errors, paradoxically leading to underestimation of trait variance attributed to shared environmental influences, and overestimation for genetic and non-shared environmental influences. Despite their minor methodological impact, parent-environment interactions should be taken seriously. For example, Hart and colleagues outline how mistaking a genetic confound for an environmental influence can inspire harmful policy [7].

As expected, increases in measurement error drove up the variance attributed to non-shared environments. There did not appear to be a bias in reduction of variance attributed to genetic or shared environmental influences. It is important to note that is not a small effect: In a trait measured with an instrument of very high reliability (α=0.9), one would expect 10% of variance to be attributable to measurement error. On averages across all simulations, even this resulted in an overestimation of non-shared environmental influences of 14 percentage points (Figure 1).

An assumption underlying twin studies that remained untested here, is that monozygotic twins are genetically identical. While this assumption does not hold [16–20], the impacts of (epi)genetical differences are hard to predict, and likely differ between traits. Hence, the conservative approach taken here was to model monozygotic twins as genetically identical.

One might wonder whether twin studies should ever be trusted. Arguably, even with increasingly accessible methods for whole-genome sequencing and the computation of polygenic risk scores, twin studies can add value beyond simple heritability questions [21]. Furthermore, many researchers are clearly diligent when conducting twin studies, honest about assumptions and limitations in their writing (e.g. [13]), and continuously looking to improve methodology (e.g. [7,15]). In sum, questions of trait heritability versus socialisation cannot be settled by twin studies alone [6], but twin studies can help answer other questions [21].

Despite all this, papers continue to be published with suspiciously high heritability estimates without plausible biological mechanisms. One famous example on the supposed genetics of political attitudes [8] is frequently cited despite thorough critiques [3,4]. A less egregious example is the “quantitative genetics of disgust” [14,22], offering seemingly sensible heritability and nonshared environment estimates of 34 and 66%, respectively [14]. However, the same estimations are yielded from a simulation with ground truths varying between 10-60 and 10-50%, respectively. While the researchers listed potential confounds, their relatively strong conclusion (doing away with the notion that early familial environments partly shape disgust) suggests that they underestimated the potential extent of these confounds. In light of the reported biases in twin studies, evolutionary psychologists would be wise to avoid employing them in the pursuit of presumed biological evidence. This is especially pertinent if no plausible mechanisms or convergent genetic evidence exist. Perhaps instead they could turn towards promising theories of cultural transmission [9–11].

In conclusion, twin studies overestimate genetic heritability in the presence of gene-environment interactions. To avoid mistaking nurture for nature, it is highly advisable that researchers measure individual and environmental variables. This will allow them to not only explicitly model parental genetic confounds (see e.g. [7]), but also direct genetic influences from twins on their environment (see e.g. [15]). While this approach will help to some degree, it is unlikely that all gene-environment interactions can be measured and modelled, and hence estimates of heritability and (the lack of) environmental influences should not be inferred from twin studies without convergent evidence from less biased designs.

## Methods

### Data simulation

Individuals’ traits were simulated using a mixture modelling approach [23], as a combination of genetic, and shared and non-shared environmental influences. Each determined a proportion of the total trait variance, which was always set to 1.

Parental polygenic traits were constructed by randomly selecting two out of 15 possible alleles for each of 10 genes, chosen so that the trait was centred on 0 with variance 1. Twin genotypes were then formed through simulated sexual reproduction with monozygotic twins inheriting the same genome, and dizygotic twins sharing ~50%. The final genetic trait (*a*) was then computed as the average of all alleles within an individual. Both shared (*c*) and non-shared (*e*) environment were sampled from normal distributions with mean 0 and variance 1. The shared environment was the same within a twin pair.

At this point, *a*, *c*, and *e* are each normal distributions centred on 0 with variance 1. Their variances were weighted by simulated proportions to make up trait *y*. The proportion of variance attributed to each component was varied between 0 and 1 in steps of 0.1; in combinations that ensured trait variance σ^2^_y_ was 1.

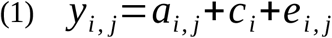

where *y_i,j_* is the trait for sibling *j* in twin pair *i*; and *a*, *c*, and *e* are normally distributed around 0 with variances σ^2^_a_, σ^2^_c_, and σ^2^_e_, respectively.

Statistically confounding the environmental influences was done by making their variances partially dependent on the genetics of twins and/or their parents. For each parent *p* of twin pair *i*, the genetic trait *a_i,p_* component was computed as the average of all alleles (same as for twins), and their combined influence on twins’ shared environment was computed as the average of both parents’ genetic trait. Parents were assumed to only impact twins’ shared environment, whereas twins were assumed to impact their shared, non-shared, or both environments. The proportions of variance in confounded shared environment *c*’ (Equation 2) and confounded non-shared environment *e*’ (Equation 3) attributed to confounders were varied between 0 and 1, in combinations that ensured component variance always added up to 1. After this, trait variance was computed in the same way as when no confounders were present (Equation 1).

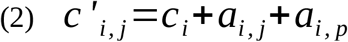

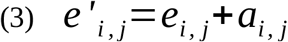

where *c’_i,j_* is the confounded shared environmental trait component for sibling *j* in twin pair *i, e’_i,j_* is the confounded non-shared environmental trait component for sibling *j* in twin pair *i*, a_i,j_ is the genetic component for sibling *j* in pair *i*, and a_i,p_ is the genetic component for both parents of sibling pair *i*. The variances of all components are set so that *c*’ and *e*’ are both normally distributed around 0, with variance 1.

### Data analysis

Generated data was analysed using traditional methods ([24], Equations 4–6) and the more contemporary approach of structural equation modelling ([25], Figure S1). Both were able to correctly identify the proportions of variance attributable to genetic heritability (*A*), and shared (*C*) and non-shared (*E*) environmental influences when no confounds were present (Figure S2).

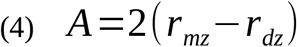

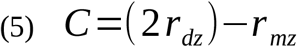

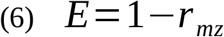

where *r_mz_* is the correlation in trait *y* between monozygotic twins, and *r_dz_* the correlation for dizygotic twins.

A total of 66 different combinations of ground truth heritability and environmental influences were simulated, in 100 runs each. Estimates from all converged models were then subtracted from the ground truth to compute the estimation error, which was then tested against the expected mean of 0 using one-sample t-tests with degrees of freedom ranging from 5770 to 6599, depending on the number of models that converged.

### Code availability

All simulation code was written in Python (v. 2.7.12; for a tutorial, see [26]), using NumPy (1.16.6) and SciPy (1.2.3) [27]; and Matplotlib (2.1.2) for generating figures [28]. Structural equation modelling was done with Lavaan (0.6-1) [29], incorporated in Python using RPy2 (2.8.6) [30]. All code is available through GitHub: github.com/esdalmaijer/2020_twin_sim

## Supplementary Figures

**Figure S1.**
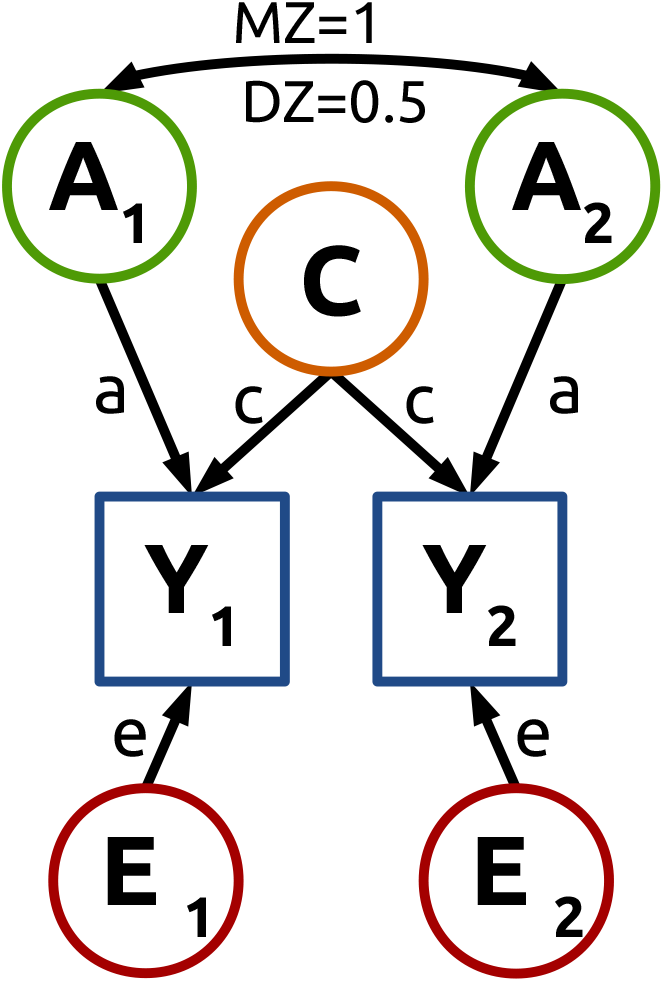
Visual representation of the structural equation model with which simulated data was analysed to estimate heritability and environmental influences.

**Figure S2.**
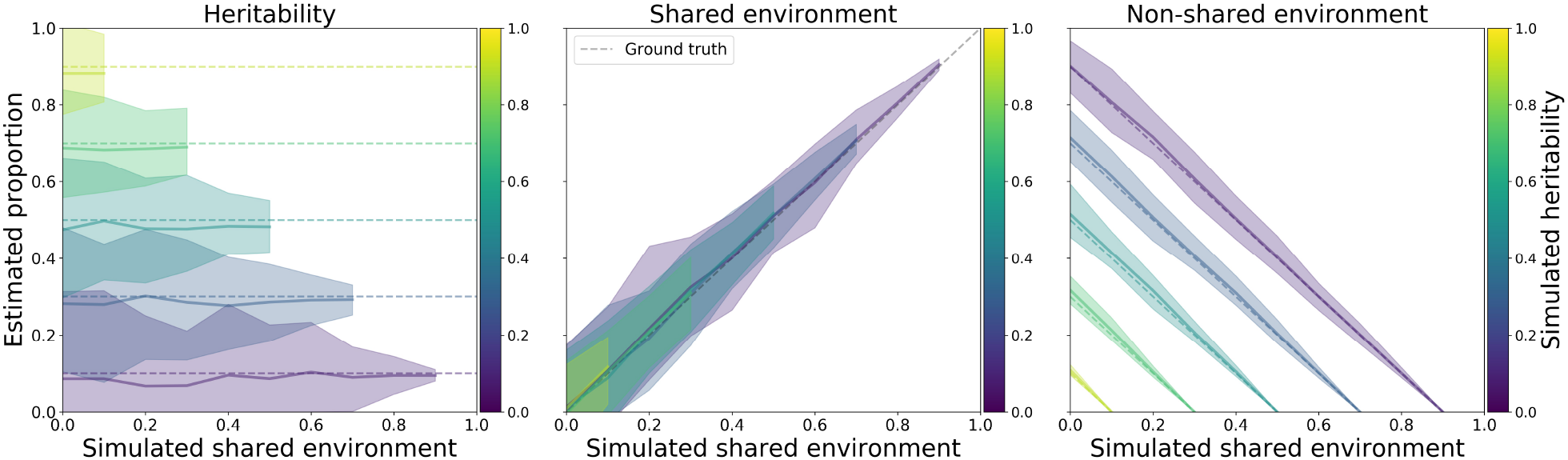
Results from 100 simulations per combination of ground truth proportions. Average estimated heritability (left panel), shared (middle panel), and non-shared environmental influences (right panel). The shaded areas indicate the 5^th^ to 95^th^ percentiles. Dashed lines show the ground truth.

**Figure S3.**
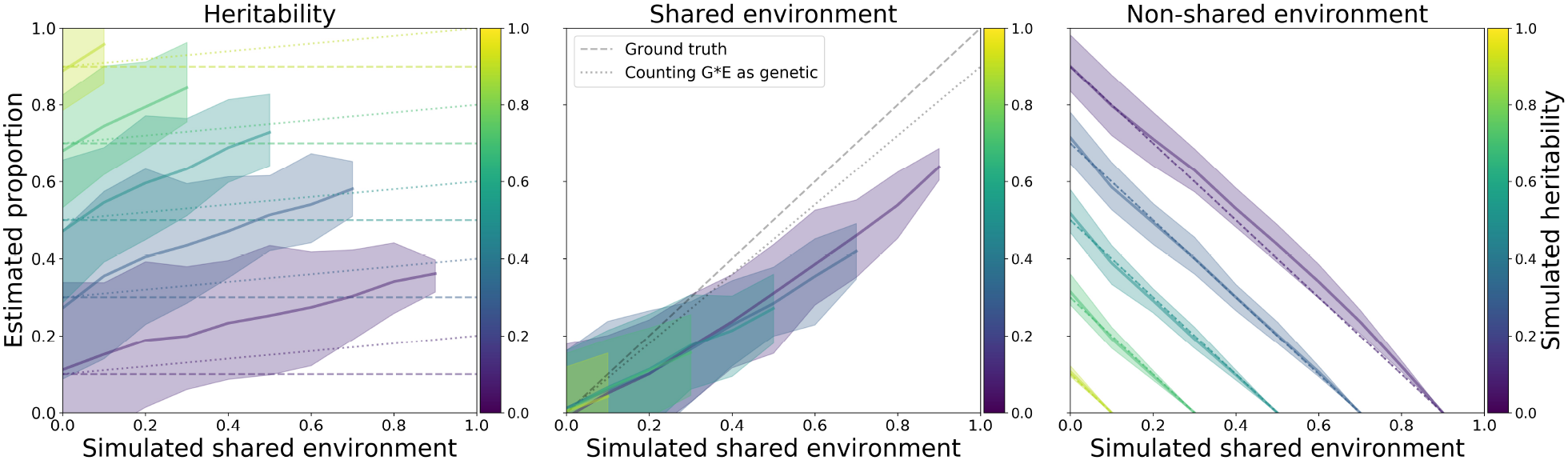
Results from 100 simulations per combination of ground truth proportions in which only shared environment was 10% confounded by twin genetics. Average estimated heritability (left panel), shared (middle panel), and non-shared environmental influences (right panel). The shaded areas indicate the 5^th^ to 95^th^ percentiles. Dashed lines show the ground truth, and dotted lines ground truth if gene-environment interactions are counted towards heritability.

**Figure S4.**
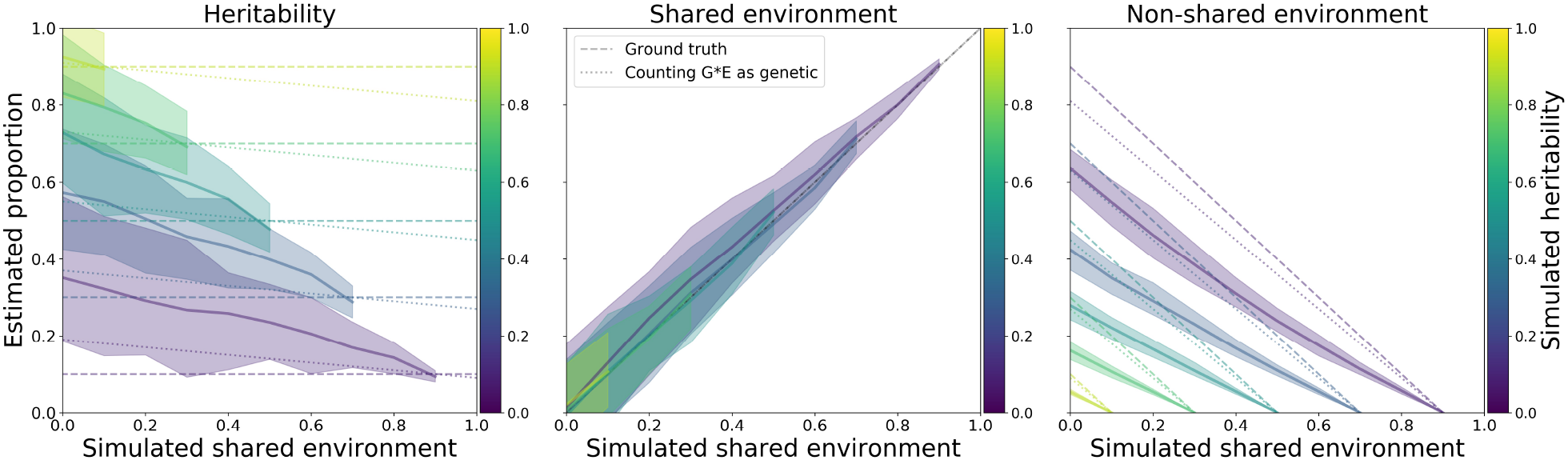
Results from 100 simulations per combination of ground truth proportions in which only non-shared environment was 10% confounded by twin genetics. Average estimated heritability (left panel), shared (middle panel), and non-shared environmental influences (right panel). The shaded areas indicate the 5^th^ to 95^th^ percentiles. Dashed lines show the ground truth, and dotted lines ground truth if gene-environment interactions are counted towards heritability.

